# ProtHyena: A fast and efficient foundation protein language model at single amino acid Resolution

**DOI:** 10.1101/2024.01.18.576206

**Authors:** Yiming Zhang, Manabu Okumura

## Abstract

The emergence of self-supervised deep language models has revolutionized natural language processing tasks and has recently extended its applications to biological sequence analysis. Traditional models, primarily based on the Transformer and BERT architectures, demonstrate substantial effectiveness in various applications. However, these models are inherently constrained by the attention mechanism’s quadratic computational complexity *O*(*L*^2^), limiting their efficiency and the length of context they can process. Addressing these limitations, we introduce **ProtHyena**, a novel approach that leverages the Hyena operator. This innovative methodology circumvents the constraints imposed by attention mechanisms, thereby reducing the time complexity to a subquadratic, enabling the modeling of extra-long protein sequences at the single amino acid level without the need to compress data. ProtHyena is able to achieve, and in many cases exceed, state-of-the-art results in various downstream tasks with only 10% of the parameters typically required by attention-based models. The architecture of ProtHyena presents a highly efficient solution for training protein predictors, offering a promising avenue for fast and efficient analysis of biological sequences.

## Introduction

Proteins, fundamental to the functionality of organisms, play diverse roles in cellular processes, ranging from biochemical reactions catalyzed by enzymes to maintaining cell shape through structural proteins. In humans, proteins supply vital amino acids that our bodies cannot synthesize independently. Understanding proteins is thus crucial to comprehending human biology and health, emphasizing the need for advanced protein representation modeling using machine learning techniques. Despite the exponential growth of protein databases in recent decades, the challenge of obtaining meaningful annotations for these sequences remains a significant barrier. The majority of proteins in these databases lack functional and structural annotations, highlighting the necessity for efficient analysis methods that can capitalize on the wealth of unlabeled protein sequences.

The adoption of self-supervised pre-training approaches from natural language processing (NLP), such as BERT [13] and RoBERTa [22], has revolutionized the representation learning of protein sequences. This methodology involves pre-training large deep learning models on millions of unlabeled protein sequences to learn universal embeddings, which are then fine-tuned for various specific protein tasks. This approach parallels the revolutionary impact of large Transformer [40] models in diverse domains like language, vision, audio, and biology [14, 29, 10]. The success of these models, particularly due to the attention mechanism, hinges on their ability to scale and facilitate in-context learning, enabling generalization to unseen data and tasks. However, a significant limitation of these models is the quadratic computational cost associated with the length of input sequences. This cost constraint severely limits the contextual capacity of the models and impedes their application to longer sequences.

In addressing the computational demands of Transformer-based models, various strategies have been employed, such as linearized, low-rank, and sparse approximations [8, 41, 39]. While these methods effectively reduce the computational load, they often necessitate a compromise between expressivity and processing speed. Hybrid models that blend these approximations with standard attention layers are proposed to strike a balance close to the performance of conventional Transformers. Foundation models (FMs) like Protein RoBERTa [16] have pioneered the use of Longformer [5] to process up to 2,048 tokens for protein sequence inputs. Similarly, scBERT [43] leverages the Performer [9] for analyzing large single-cell gene expression data, and MolFormer [35] incorporates linear attention [19] to more effectively capture spatial relations between chemical atoms.

Despite these advancements, a significant challenge persists with the size of these models, often exceeding 100 million parameters. Most protein-related foundation models [26, 16, 7, 42] are limited to handling inputs of 512 tokens. Moreover, they adopt byte pair encoding (BPE) [37] as a method for representing protein sequences, analogous to tokenizing words in language processing. This approach aggregates meaningful protein units, but it faces limitations in the context of single amino acid changes. The role of individual amino acids is critical, as they represent physical analogs with significant impacts. For instance, a single amino acid alteration can drastically alter a protein’s function or structure. In contrast to natural language, where the semantics can often withstand single character or word alterations over extended contexts, the domain of protein requires a nuanced approach. Proteins demand an analysis that can maintain long-range contextual understanding while simultaneously resolving single amino acid details. This requirement for both extensive context and high-resolution at the amino acid level presents a unique and substantial challenge in the field of protein, pushing the boundaries of current machine learning models and demanding innovative solutions to accurately model these complex biological sequences.

The recent development of Hyena [28], a large language model based on implicit convolutions, marks a significant advancement in the field of machine learning. Hyena has demonstrated its ability to match the quality of attention-based models while significantly reducing computational time complexity. This efficiency enables the processing of longer contextual sequences. Motivated by these advancements, we present ProtHyena, a fast and parameter-efficient foundation model that incorporates the Hyena operator for the analysis of protein data. This architecture can unlock the potential to capture both the long-range and single amino acid resolution of real protein sequences over attention-based approaches.

To assess the efficacy of ProtHyena, we initially pretrain the model using the Pfam [17] dataset and then fine-tune it for a variety of tasks. For the first time, we have fine-tuned our model across 10 distinct protein-related downstream tasks and have documented their respective performances. The results demonstrate that ProtHyena approaches or even surpasses state-of-the-art performance in some of these tasks. In the spirit of facilitating comprehensive evaluations for future researchers working on novel protein language models, we have made these datasets publicly available. This initiative aims to provide a more thorough framework for assessing the capabilities of emerging models in the field of protein sequence analysis.

## Background

### Self-Attention

The self-attention operator, a key component of Transformer models as introduced by [40], plays a pivotal role in processing sequences. When given a sequence *x* of length *L* and dimension *D*, each head of the self-attention mechanism transforms *x* into an output *y* using the self-attention operator *A*(*x*):

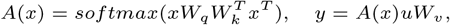

where *W*_*q*_, *W*_*k*_, and *W*_*v*_ are learnable linear projections for the query, key, and value, respectively. The softmax and scaling function, denoted as *softmax*, is also applied. This mechanism allows the model to capture pairwise relationships among all tokens in the sequence, thereby understanding the global context.

However, a notable drawback of self-attention is its high computational cost for long sequences, with complexity scaling quadratically with the sequence length *O*(*L*_2_ ). To counteract this, various methods have been developed. One such method is factorized self-attention, used in sparse Transformers. This technique reduces memory and computational demands by allowing self-attention heads to focus on a subset of tokens. Another method is the Performer, which addresses the memory complexity of self-attention by decomposing the self-attention matrix. This allows the Performer to implicitly store the attention matrix with linear memory complexity, thereby enabling the processing of longer sequences than traditional Transformers. As an alternative, linear attention methods create approximations of the self-attention operator *A*(*x*) that can be computed more quickly, with less than quadratic time complexity. However, while these methods allow for processing longer sequences due to their reduced time complexity, they often come with a trade-off, resulting in a decrease in the model’s expressivity.

To match the high performance of Transformers with a lower computational burden, it’s crucial to obtain operators that possess attention’s three defining properties: data control, sublinear parameter scaling, and unrestricted context. [28] successfully achieves this by introducing an operator characterized by a recurrent structure, which combines two subquadratic primitives: a long convolution and an element-wise multiplicative gating. This innovative operator retains the core attributes of attention, ensuring efficient processing across lengthy sequences without compromising on the ability to model complex dependencies.

### Large Convolutional Models and Hyena Operator

Inspired by discrete-time systems in signal processing, the Linear State-Space Layers (LSSL) model [18] maps sequences by simulating a linear discrete-time state-space representation.

A discrete convolution between an input *x* of length *L* and a learnable filter *h* is defined as follows:

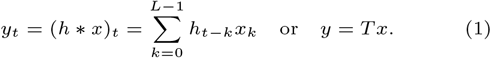

where *T* in ℝ^*L×L*^ is the Toeplitz matrix corresponding to the convolution. In the realm of deep learning and signal processing, convolutions have historically been pivotal. The Toeplitz matrix *T* in ℝ^*L×L*^ is foundational to convolution operations. Recent advancements demonstrate that stacking a series of long convolution layers, where the number of layers *τ* is determined by a specific function of the sequence length *L, τ* := *γ*_*θ*_(*L*), leads to cutting-edge results across benchmarks involving long sequences. This approach has been refined through various methodologies like state-space models and implicit parametrizations using neural fields. To better model language, H3 [12] and Hyena [28] have leveraged implict long convolutions and gating mechanisms to attain Transformer-level efficiency within a time complexity of *O*(*L* log_2_ *L*), which is significantly more efficient than the *O*(*L*_2_ ) time complexity of attention-based models.

ProtHyena is inspired by these advancements, demonstrating that models capable of processing long-contexts without relying on attention can achieve high performance in protein tasks. This approach extends the range of protein sequence analysis, opening up new possibilities such as in-context learning.

## Methods

### Tokenization

In our approach, we use the natural protein vocabulary, treating each amino acid as an individual token. These tokens encompass the 20 standard amino acids, represented by the characters ‘D’, ‘N’, ‘E’, ‘K’, ‘V’, ‘Y’, ‘A’, ‘Q’, ‘M’, ‘I’, ‘T’, ‘L’, ‘R’, ‘F’, ‘G’, ‘C’, ‘S’, ‘P’, ‘H’, and ‘W’. Additionally, we include tokens for special cases: ‘X’ for any or unknown amino acids, ‘U’ for Selenocysteine, ‘B’ representing either Asparagine or Aspartic acid, ‘O’ for Pyrrolysine, and ‘Z’ for either Glutamic acid or Glutamine. Beyond these, our model also incorporates special character tokens to signify padding, separation, and unknown characters. Each of these tokens is mapped to an embedding with dimension D, facilitating the representation and processing of protein sequences within our model framework. This method allows for a comprehensive and accurate encoding of protein sequences, crucial for effective analysis and modeling.

To provide a comparative analysis, we also trained a variant of our model using byte pair encoding (BPE) [37], named ProtHyena-bpe. This model employs a BPE tokenizer with a vocabulary size of 10k, which consequentially expands the model parameters to 4.3 million. This setup allows us to directly compare the performance of a model with single amino acid resolution against one that utilizes data compression through BPE tokenization, shedding light on the trade-offs and benefits associated with each method.

### ProtHyena Framework

The overall framework of our ProtHyena framework is shown in Fig 2.

The Hyena operator is defined by a recurrent structure, incorporating long convolutions and element-wise gating, as shown in the equation:

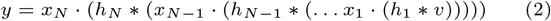

where *v, x*_1_, …, *x*_*N*_ denote the input projections, *N* represents the number of recurrences, · indicates the element-wise gating and *\** denotes the long convolution.

To be more specific, Fig.1 provides a visual representation of the Hyena operator with *N* = 2. When processing an input sequence *x* of length *L*, the Hyena operator is applied as follows:

**Fig. 1.**
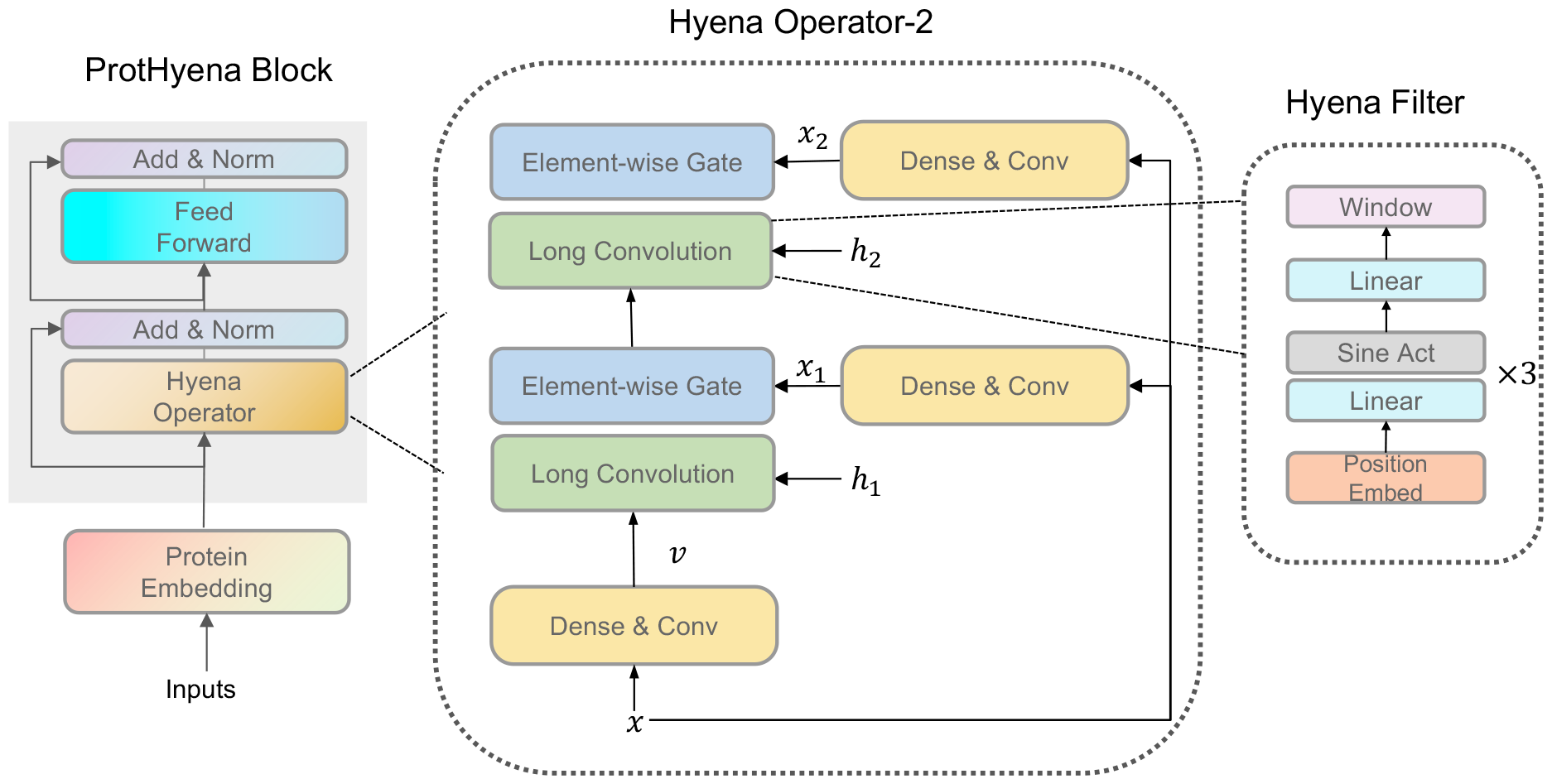
The architecture of the ProtHyena block with order=2. At its core is the Hyena operator, which utilizes extended convolutions coupled with element-wise gating mechanisms. The input undergoes a transformation through dense layers and short convolutions to feed into the gates. These extended convolutions are implicitly parameterized by an MLP (multilayer perceptron), responsible for generating the convolutional filters. The convolution process itself is efficiently executed using a Fast Fourier Transform (FFT), ensuring a lower time complexity of *O*(*L* log_2_ *L*), which is crucial for managing long sequences effectively.

**Fig. 2.**
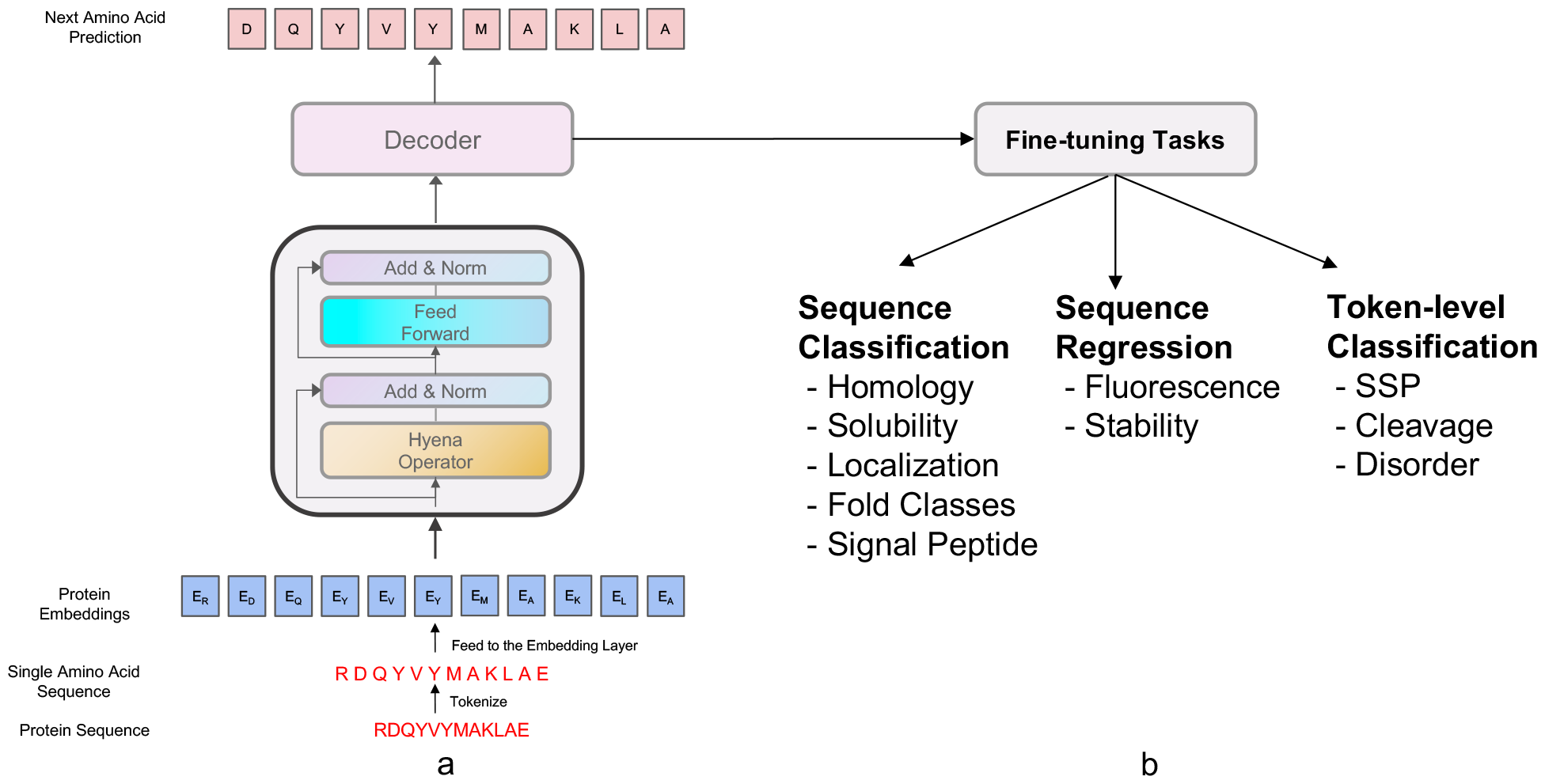
Overview of our proposed framework for ProtHyena. (a) the Generative Pre-training strategy. In this phase, the input protein sequence is tokenized, with each amino acid represented as a token. The model structure includes two layers of Hyena Blocks; each layer consists of a Hyena Operator and a fully connected network, supplemented by residual connections and layer normalization. The sequence is then passed through a decoder which predicts the next amino acid in the sequence. (b) the fine-tuning process applied to the pre-trained model across ten different downstream tasks. These tasks consists a range of protein sequence analyses, including Sequence Classification for Homology, Solubility, Localization, Fold Classes, and Signal Peptide, Sequence Regression for tasks like Fluorescence and Stability, and Token-level Classification tasks such as SSP, Cleavage, and Disorder. The model is individually fine-tuned for each task to optimize its predictive performance on that specific task.

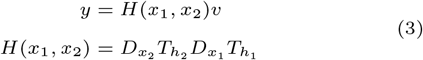

where (*x*_1_, *x*_2_, *v*) are projections of the inputs, specifically generated by a linear projection followed by 1 dimension short convolution. The Toeplitz matrix 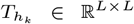*∈ ℝ*^*L×L*^ is created from a learnable long convolution filter generated as the output of a neural network, with each element 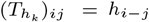. The values of the convolution filter are derived from a small neural network, denoted as *γ*(*θ*), which takes the time index and protein embeddings as its input. This function, expressed as *h*_*t*_ = *γ*(*θ*(*t*)), allows the operator to efficiently handle sequences without a linearly increasing in the number of parameters. This setup ensures that our model remains parameter-efficient even as it scales to accommodate very long sequences. Moreover, the diagnoal matrices 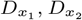 *∈* ℝ^*L×L*^ act as gates, operating element-wise on the projections. These projections are generated through the application of a dense linear layer followed by short convolutions on the input sequence. Upon obtaining the filter values and projecting the input signal, we utilize the Fast Fourier Transformation (FFT) method, which operates at a computational complexity of *O*(*L* log_2_ *L*) in the frequency domain.

### Pre-training

The ProtHyena model embodies a decoder-only, standard sequence-to-sequence Generative Pre-Training (GPT) [30] procudure. The GPT architecture is originally a language model designed for improving language understanding by predicting word adapted here for protein sequence analysis. Its pre-training involves unsupervised learning from extensive sequence data to learn the statistical properties of amino acids in proteins. GPT employs an autoregressive framework, which, during pre-training, maximizes the probability of the next amino acid given the preceding sequence of amino acids.

The objective of the GPT model is to maximize the joint probability of a sequence of amino acids. Given a sequence of protein consists of amino acids *a*_1_, *a*_2_, …, *a*_*N*_, the model aims to maximize the likelihood of the sequence *P* (*a*_1_, *a*_2_, …, *a*_*N*_ ). This is achieved through the following autoregressive formula:

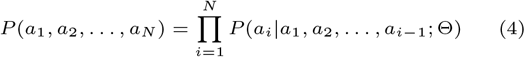

where *P* (*a*_*i*_|*a*_1_, *a*_2_, …, *a*_*i*−1_ ) represents the conditional probability of amino acid *a*_*i*_ given all preceding amino acids in a specific protein sequence (*a*_1_, *a*_2_, …, *a*_*i*−1_ ). The conditional probabilities are parameterized by the transformer model in conventional GPT model frameworks. In our current work, we have adopted the Hyena architecture to parameterize these probabilities. This architectural choice reflects our aim to utilize the Hyena’s efficient handling of longer sequence contexts, which is crucial for modeling the intricacies of sequential data more effectively than traditional transformer models.

During pre-training, GPT aims to minimize the cross-entropy loss of the next amino acid prediction. For each amino acid *a*_*i*_, the model generates a probability distribution 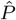 and compares it with the actual next amino acid *a*_*i*+1_ in a one-hot encoded format. The loss function is defined as:

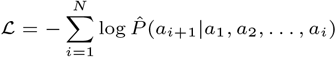

The GPT procedure adopts an autoregressive framework originally based on the Transformer architecture, adapted for protein sequence analysis. Its pre-training objective is to capture the intrinsic structure of protein sequences by maximizing the joint probability distribution of these sequences. Optimization is achieved by minimizing the crossentropy loss, which measures the discrepancy between the model’s predicted amino acid distribution and the actual distribution.

## Experiments

In this section, we evaluate our ProtHyena model by pretraining and fine-tuning on different protein analysis tasks. We first introduce the pre-training settings, then the finetuning tasks, and report the results on these different protein downstream tasks

### Pre-training data

Consistent with prior studies [31, 32], our pre-training utilizes the Pfam database [24], a comprehensive dataset encompassing 14 million protein domains commonly employed in the field of Bioinformatics. This dataset groups protein sequences into families based on evolutionary relationships. We have split the dataset at random into separate sets for training, validation, and test. As a measure of our pre-training effectiveness, we report the model’s perplexity.

Perplexity is a measurement used in the field of natural language processing and, by extension, in protein language modeling, to evaluate the probability model’s ability to predict a sequence. It gauges how well a probability distribution or probability model predicts a sample. A lower perplexity score indicates that the model predicts the sample more accurately, implying a better performance. It is mathematically defined as the exponentiated average negative log-likelihood of a protein sequence. If we consider a protein sequence of length *L*, with *W* representing the sequence of tokens (amino acids in the case of proteins), the perplexity (*PPL*) is computed using the following formula:

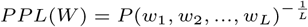

where *P* (*w*_1_, *w*_2_, …, *w*_*L*_) is the probability of the sequence according to the model. This can be further broken down into:

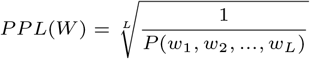

In our case, for numerical stability and to simplify the computation, perplexity is calculated as the exponentiation of the cross-entropy loss:

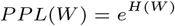

where *H*(*W* ) is the cross-entropy of the sequence *W*, defined as:

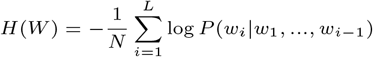

A lower perplexity score indicates a protein language model’s enhanced learning of protein sequences.

### Fine-tuning downstream tasks

To evaluate ProtHyena, we have consolidated the assessment datasets from prior protein language modeling efforts and, to the best of our knowledge, conducted the most extensive performance testing on a diverse array of downstream tasks to date. ProtHyena was evaluated on ten benchmarks that span the core areas of protein research. These benchmarks include a wide range of protein-related aspects such as function, structure, post-translational modifications, and biophysical properties, as detailed in Table 1.

**Table 1.** Summary of downstream fine-tuning tasks. For multiclass classification tasks, the number of classes is in the parentheses.

| Task | Type | Resolution | Metric | Source |
| --- | --- | --- | --- | --- |
| Secondary Structure | Multi-class(3) | Token | Accuracy | [25, 31] |
| Disorder | Binary | Token | Accuracy | [25] |
| Remote Homology | Multi-class(1195) | Sequence | Accuracy | [3, 4, 31] |
| Localization | Multi-class(10) | Sequence | Accuracy | [1] |
| Fold Classes | Multi-class(7) | Sequence | Accuracy | [3, 4] |
| Signal Peptide | Binary | Sequence | Accuracy | [2] |
| Neuropeptide Cleavage | Binary | Token | Accuracy | [6] |
| Solubility | Binary | Sequence | Accuracy | [20, 16] |
| Fluorescence | Regression | Sequence | Spearman’s $\rho$ | [36, 31] |
| Stability | Regression | Sequence | Spearman’s $\rho$ | [34, 31] |

### Pre-training and fine-tuning settings

For pre-training our protein models, we adopted the base configuration as per the guidelines in [27]. Our setup starts with 2 Hyena layers with a recursion order of N=2. The model’s embedding size is set to 256, and it contains 1024 feed-forward hidden units. To effectively demonstrate the Hyena operator’s capabilities, we also trained two decoder-only transformer models, named ProtGPT-tiny and ProtGPT-base, for comparison. ProtGPT-tiny is composed of 2 transformer decoder layers. It matches ProtHyena in terms of embedding size and feed-forward hidden units, equating to an identical count of trainable parameters. In contrast, ProtGPT-base is more extensive, with 8 transformer decoder layers, an embedding size of 512, and 2048 feed-forward hidden units. We managed our batch sizes to 256, and during training, we maintained a maximum protein sequence length of 1024 for ProtHyena and ProtGPT-tiny. Due to memory constraints, the maximum length for ProtGPT-base was limited to 512. Training was performed utilizing the Adam optimizer [23], starting with an initial learning rate of 0.0006 and employing a cosine decay learning schedule. The total number of training steps was approximately 40k. Throughout pre-training, we used perplexity as our primary performance metric.

## Main Results

### ProtHyena performs better than Attention based model on protein modeling

In our experiments, ProtHyena demonstrated superior performance in protein modeling compared to attention-based methods. To evaluate the effectiveness of models based on the Hyena Operator, we trained two additional models: ProtGPT-tiny, with a parameter size of 1.6 million, matching that of ProtHyena, and the larger ProtGPT-base, encompassing 6.6 million parameters. Both models underwent training on the same Pfam dataset as ProtHyena, using a batch size of 256 and for an equal number of steps.

The test results, as presented in Table 2, reveal that ProtHyena achieved the lowest perplexity among the models, indicating its enhanced performance. This outcome underscores the effectiveness of ProtHyena in protein modeling, outperforming its counterparts in handling the complexities of protein sequences. The lower perplexity of ProtHyena, particularly in comparison to the larger ProtGPT-base, highlights its efficiency and the robustness of the Hyena Operator in modeling proteins, validating our model’s superior design and capabilities in this domain.

**Table 2.** Perplexity on Pfam test set. Lower is better.

| Model | PERPLEXITY | Model Size (M) |
| --- | --- | --- |
| ProtGPT-tiny | 2.478 | 1.6 |
| ProtGPT-base | 2.000 | 25.2 |
| ProtHyena-bpe | 1.469 | 4.3 |
| ProtHyena | 1.466 | 1.6 |

### ProtHyena can model ultra-long protein sequence faster than Transformers

ProtHyena demonstrates notable advantages in terms of speed and the ability to model longer sequences compared to attention-based models. In our benchmark tests, we compared the runtime of an order 2 Hyena operator against both traditional attention and flash-attention layers [11]. The Hyena operator employs a fused CUDA kernel for executing FFTConv operations. We set the batch size to 1 and measured the runtime in milliseconds, with the results displayed in Fig 3.

**Fig. 3.**
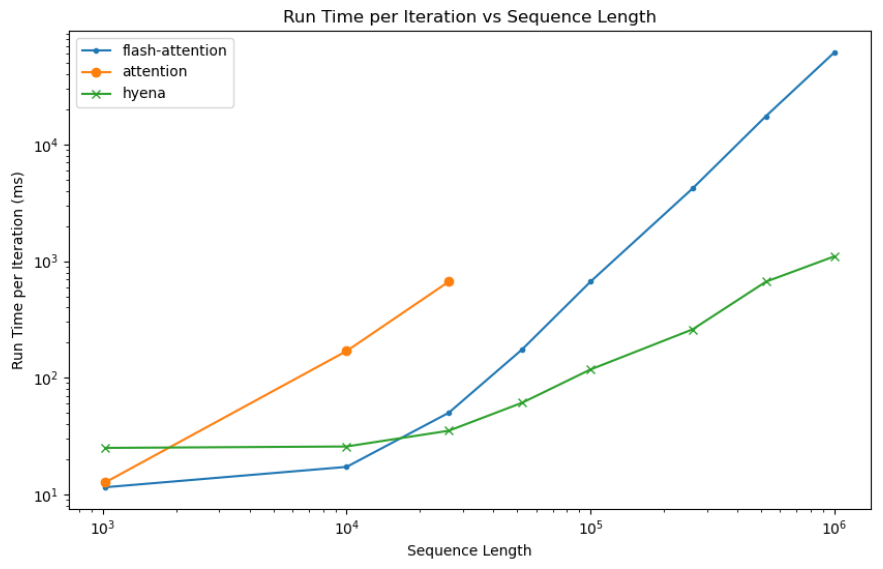
Benchmarks the runtime performance of the hyena, attention, and flash-attention mechanisms across different sequence lengths on a single 80G A100 GPU with both the x and y axes plotted on a log scale. Each model was tested with batch size 1. Notably, the Attention mechanism’s plot line has gaps for sequence lengths exceeding 20,000 due to CUDA out-of-memory errors.

Our findings show that Hyena achieves speedups of up to 60 times at a sequence length of 1M. The crossover point, where Hyena starts outperforming traditional attention, is at a sequence length of 2048, and for flash-attention, it lies between 10k and 20k. Notably, when the sequence length exceeds a certain threshold, standard attention methods lead to CUDA memory exhaustion on an 80G A100 GPU, and flash-attention also significantly slows down.

Despite Hyena’s absolute reduction in runtime, its speed advantage becomes more evident only with longer sequences where the difference in processing time becomes significantly large. This is because Hyena’s hardware utilization is lower than that of flash-attention. However, we anticipate the gap between the theoretical maximum speedup and actual performance to decrease with more optimized implementations of FFTConv and advancements in specialized hardware.

To process long protein sequences, previous methods primarily employed frequency-based byte pair encoding (BPE), which aggregates proteins into larger, supposedly meaningful units, often truncating them to a length of around 512. This approach, while useful, results in the creation of extensive new vocabularies that are considerably larger than the natural vocabulary of just 20 amino acid (plus 5 unknown/non-standard amino acids). As noted by [38], these expanded vocabularies tend to be less generalizable, and the truncation process can lead to the loss of crucial protein information.

In contrast, our model, ProtHyena, introduces a significant breakthrough by being capable of modeling protein sequences up to 1 million amino acids in length. This capability far exceeds the limits of previous methods, addressing the key challenges of information loss due to sequence truncation and the limitations imposed by expanded vocabularies. ProtHyena’s capacity to handle such extraordinarily long sequences opens new horizons in protein sequence analysis, allowing for a more comprehensive and uninterrupted understanding of protein structures and functions.

Furthermore, the remarkable speedup achieved by ProtHyena, reaching up to 60 times faster than conventional models at longer sequence lengths, is a testament to its optimized architecture. The efficiency gains observed in our benchmarks, especially at crossover points with traditional attention and flash-attention, highlight the model’s superior design in balancing computational load and performance.

### ProtHyena performance on downstream tasks

#### Results on TAPE benchmark

In this section, we evaluated the newly developed ProtHyena method on the TAPE test dataset against an array of contemporary state-of-the-art algorithms including BERT Transformer and LSTM models in TAPE [31], ProtT5 [15], ProteinBERT [7], and SPRoBERTa [42]. The ‘contact prediction’ task from the TAPE benchmark was excluded from this analysis because it is incompatible with the model’s output format. ProtHyena is designed to generate sequence-level and token-level predictions, and it does not provide pairwise predictions required for contact prediction tasks. The experimental results of different methods are shown in Table 3.For Secondary Structure Prediction (SSP) and Remote Homology, accuracy is the chosen metric, while for Fluorescence and Stability, performance is gauged using Spearman’s *ρ*.ProtHyena stands out in Remote Homology with the highest accuracy of 0.317, indicating a significant improvement over other models like TAPE Transformer and SPRoBERTa, which score 0.210 and 0.230, respectively. In terms of Fluorescence, ProtHyena also leads with a Spearman’s *ρ* of 0.678, showcasing its robustness in capturing complex protein properties.

**Table 3.**
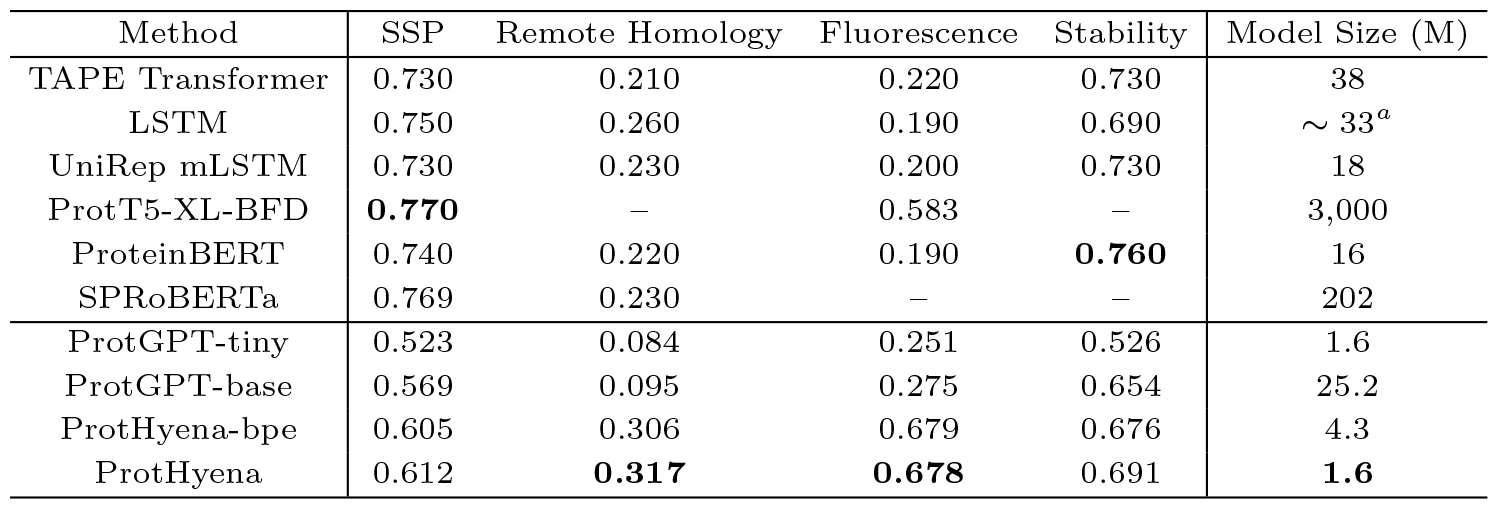
Evaluation on TAPE benchmark datasets. Main results of our ProtHyena model and other compared methods. ProtGPT-tiny and PortGPT-base is our reproduced backbone model that trained only on the amino acid representation. The results for these baseline methods are reported from the original paper, and ‘-’ means this task is not reported in the original paper

| Method | SSP | Remote Homology | Fluorescence | Stability | Model Size (M) |
| --- | --- | --- | --- | --- | --- |
| TAPE Transformer | 0.730 | 0.210 | 0.220 | 0.730 | 38 |
| LSTM | 0.750 | 0.260 | 0.190 | 0.690 | ~ 33<sup>a</sup> |
| UniRep mLSTM | 0.730 | 0.230 | 0.200 | 0.730 | 18 |
| ProtT5-XL-BFD | **0.770** | – | 0.583 | – | 3,000 |
| ProteinBERT | 0.740 | 0.220 | 0.190 | **0.760** | 16 |
| SPRoBERTa | 0.769 | 0.230 | – | – | 202 |
| ProtGPT-tiny | 0.523 | 0.084 | 0.251 | 0.526 | 1.6 |
| ProtGPT-base | 0.569 | 0.095 | 0.275 | 0.654 | 25.2 |
| ProtHyena-bpe | 0.605 | 0.306 | 0.679 | 0.676 | 4.3 |
| ProtHyena | 0.612 | **0.317** | **0.678** | 0.691 | **1.6** |

#### Results on other downstream tasks

Tables 4 and 5 summarize the performance of various models on ProteinBERT benchmarks, including tasks like Disorder, Fold Classes, Signal Peptide, Cleavage also Localization and Solubility.

**Table 4.**
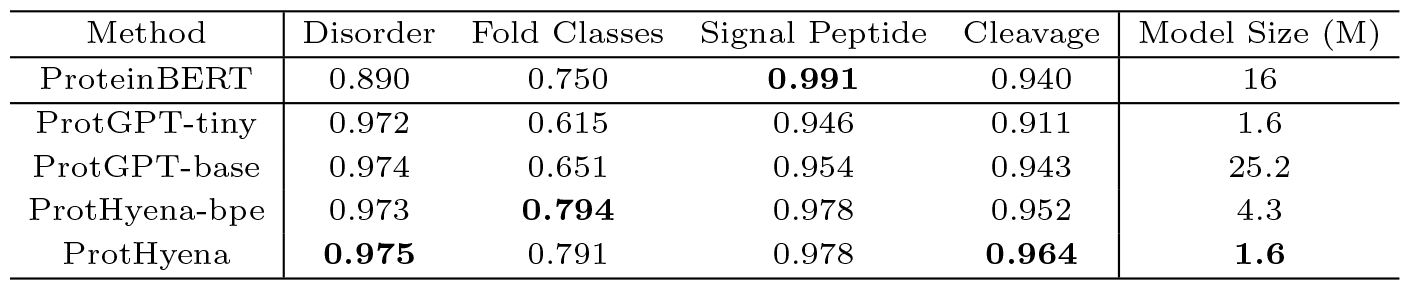
Evaluation on Other ProteinBERT benchmarks.

**Table 5.** Accuracy on Localization and Solubility tasks.

| Method | Localization | Solubility | Model Size (M) |
| --- | --- | --- | --- |
| DeepLoc | 0.780 | – | – |
| DeepSol | – | **0.770** | – |
| ProteinRoBERTa | 0.793 | 0.580 | 170 |
| ESM-1b | 0.800 | 0.673 | 650 |
| ESM-2 | 0.818 | 0.604 | 15000 |
| ProtT5-XL-U50 | **0.832** | 0.744 | 3000 |
| Ankh_base | 0.814 | 0.764 | 453 |
| ProtGPT-tiny | 0.529 | 0.686 | 1.6 |
| ProtGPT-base | 0.589 | 0.688 | 25.2 |
| ProtHyena-bpe | 0.659 | 0.729 | 4.3 |
| ProtHyena | 0.667 | 0.725 | **1.6** |

ProtHyena showcases exceptional performance, achieving the highest accuracy scores in Disorder (0.975) and Fold Classes (0.791) and competitive scores in simple Signal Peptide (0.978) and Cleavage (0.947) tasks, outperforming ProteinBERT and ProtGPT models.

For protein localization and solubility tasks, we compared against ProteinRoBERTa [16], ESM-1b [33] and ESM-2(15B) [21]. We also compared to ProtT5-XL-U50 [15] and Ankh base models. The ProtHyena demonstrates a balance of accuracy and solubility scores at 0.667 and 0.725 respectively, surpassing the ProtGPT variants and approaching the scores of specialized models like DeepSol [20].

It is noteworthy that ProtHyena achieves these results with a significantly smaller model size of only 1.6 million parameters, which is a fraction of the size of other high-performing models like ESM-1b and ESM-2, highlighting its efficiency. The modest size of ProtHyena does not compromise its accuracy, making it a highly efficient and promising model for a wide range of protein-related tasks.

In our experimental analysis, while ProtHyena-bpe exhibits slight advantages in fold classes and solubility tasks, the ProtHyena model with single amino acid level tokenization outperforms not only in token-level classification tasks but also shows superior performance across a broader range of tasks. Our findings reinforce the value of single amino acid resolution in protein language modeling, offering a detailed and nuanced understanding of protein sequences which is critical for advancing the field.

ProtHyena distinguishes itself by achieving commendable results with a model size that is remarkably lite 1.6 million parameters, which pales in comparison to the several hundred to thousandfold larger parameter counts of most state-of-the-art models. Furthermore, while these larger models are pre-trained on protein sequences ranging from hundreds to thousands of millions, ProtHyena’s training is conducted on a more modest dataset of approximately 10 million sequences. Despite this, ProtHyena stands out in half of the evaluated tasks, exhibiting good performance. This efficiency makes it a highly efficient and promising model for a wide range of protein-related tasks.

## Conclusion

In this paper, we presented ProtHyena, a novel protein language model that integrates the Hyena operator to address the computational challenges encoutered by attention-based models. ProtHyena not only efficiently processes exceptionally long protein sequences but also delivers, and sometimes outperform, state-of-the-art performance in a wide array of downstream tasks. With only a fraction of the parameters required by traditional models, ProtHyena exemplifies a significant leap forward in the field of protein sequence analysis. Our comprehensive pre-training and fine-tuning on the expansive Pfam dataset across ten different tasks have demonstrated the model’s ability to capture complex biological information with high accuracy. The adoption of the Hyena operator enables ProtHyena to perform at a subquadratic time complexity, making it a highly fast and efficient tool for analyzing biological sequences.

In the future, we aim to scale up ProtHyena to fully leverage its capabilities and precision, while also exploring masked language modeling methods for pre-training to broaden its applicability. The architecture we have introduced sets a new precedent for protein language models, offering a promising framework for future advancements in the field.

